# MAP kinase Slt2p attenuates cell wall mRNA decay by downregulating the RNA-binding protein Rbp1p in response to stress

**DOI:** 10.1101/2020.09.30.319939

**Authors:** Lin-Chun Chang, Yu-Chieh Wu, Yu-Yun Chang, Fang-Jen Lee

**Affiliations:** Institute of Molecular Medicine, College of Medicine, National Taiwan University, and Department of Medical Research, National Taiwan University Hospital, Taipei, Taiwan

**Keywords:** RNA-binding protein, posttranscriptional regulation, kinase, cell wall stress

## Abstract

The yeast cell wall integrity (CWI) MAPK pathway is a signaling cascade function in maintaining cell wall integrity under stressful environmental conditions. Recently, the activity and signaling of Slt2p (Mpk1p) MAP kinase has been shown to control assembly of the processing body (P-body) upon cell wall stresses, implicating its posttranscriptional role in decay of cell wall mRNAs. However, how Slt2p MAP kinase directly regulates the stability of cell wall transcripts during cell wall stress remains unclear. Here, we reported that the RNA-binding protein Rbp1p (Ngr1p) is a downstream effector and target of Slt2p MAP kinase during activation of the cell wall stress signaling cascade. In addition to the well-defined target mitochondrial porin mRNA, we found that Rbp1p also negatively regulates the stability of a subset of Slt2p-regulated cell wall transcripts. Deletion of *RBP1* increases the level of cell wall transcripts and partially suppresses the hypersensitivity of the *slt2Δ* deletion strain to cell wall damage. Slt2p is necessary for cell wall stress-induced stabilization of cell wall transcripts. Deletion of *RBP1* compromises the destabilization of cell wall transcripts in *slt2Δ* mutants under cell wall stress. Notably, C-terminal deleted Slt2p impairs its function in promoting turnover of the Rbp1p protein and fails to stabilize cell wall transcripts, although it can complement the growth defect of the *slt2Δ* strain upon cell wall stress. Altogether, our results demonstrate that MAP kinase Slt2p attenuates CWI mRNA decay in response to cell wall damage by downregulating the activity of the RNA-binding protein Rbp1p.

## Introduction

In *Saccharomyces cerevisiae*, the degradation of mRNAs is composed of several processes. Deadneylation of the 3’-poly (A) tail first occurs, followed by decapping of the 5’ end of mRNAs. Both processes expose the internal and vulnerable regions of mRNAs to be more accessible for 5’-3’ exonuclease Xrn1p-directed mRNA hydrolysis or 3’-5’ degradation by the Ski exosome complex [1]. Moreover, mRNA degradation is facilitated by mRNA-binding proteins (RBPs), such as Rbp1p, Pub1p and Puf family proteins, which recruit the decay machinery to mRNAs that are tagged for degradation [2,3]. Proteins involved in mRNA degradation and nontranslating mRNAs assemble into cytoplasmic foci, which are called processing bodies (P-bodies) and are strongly induced in response to stress conditions, such as glucose starvation and osmotic and heat stresses [4–6]. While mRNA degradation has been imaged throughout the cytoplasm in cells [7–9], most recent models of P-bodies function as storage granules containing translationally repressed mRNAs and inactive decay enzymes [10].

We have previously identified the RNA-binding protein Rbp1p, a protein that contains three RNA recognition motifs (RRMs) and a C-terminal Asn-Met-Pro-rich (NMP) region and negatively regulates cell growth [11]. *RBP1* is not an essential gene (systematic name: YBR212W; standard name: *NGR1* (negative growth regulator 1)) due to a slow-growth phenotype in yeast upon overexpression [12]. We found that Rbp1p binds the 3’-untranslated region (UTR) of mitochondrial porin mRNA via its RRM domains [13]. This interaction with Rbp1p accelerates the turnover of porin mRNA in an exonuclease Xrn1p-dependent manner [14]. More recently, findings suggest that Rbp1p directly binds the nonconserved C-terminal region of RNA helicase Dhh1p, a component of the mRNA decay machinery. We then proposed that Rbp1p brings mRNA decay machinery to the vicinity of porin mRNA and promotes the degradation of porin mRNA [3]. In addition to controlling the stability of porin mRNA, Rbp1p localizes to P-bodies in response to stresses [14]. The Protein Kinase A (PKA) and High Osmolarity Glycerol (HOG) MAPK pathways have been shown to regulate the assembly of P-bodies and mRNA decay in response to glucose starvation and osmotic stress, respectively [15,16]. It remains unclear whether PKA, HOG or other signaling pathways could control Rbp1p activities in mRNA decay and P-body localization upon stress.

Recently, the cell wall integrity (CWI) MAPK pathway has been reported to control P-body assembly during cell wall stress, implying its posttranscriptional role in the regulation of gene expression [17]. The CWI pathway monitors cell wall stress as well as other environmental changes and directs cell wall remodeling accordingly. Factors that activate CWI signaling range from cell wall stress agents, such as Congo Red, to unfavorable growing temperatures and osmotic shock [18]. Information on these stresses is first detected by a variety of cell membrane sensors but eventually converges to the central molecules, the Slt2p mitogen-activated protein (MAP) kinase cascade [18]. The Slt2p MAP kinase cascade includes upstream MAPKKK Bck1 and MAPKKs Mkk1/Mkk2, and the activation of Slt2p requires a series of phosphorylation events. Two transcription factors, Rlm1p and SB (Swi4p/Swi6p), work downstream of Slt2p to turn on the transcription of mRNAs, whose protein products are essential for cell wall repair or remodeling [18,19]. A genome-wide survey revealed that at least 20 functional-related mRNAs were upregulated in an Rlm1p-dependent manner [20]. Moreover, based on global transcriptional analyses, most cell wall-related mRNAs significantly increased after exposure to cell wall stresses, but this upregulation was lost in the *slt2*Δ mutant [21]. As Slt2p has indisputable roles in transcriptional reprogramming and cell wall stress [18], how cell wall integrity signaling and Slt2p MAP kinase manage cell wall stresses at the posttranscriptional level, especially the stability of cell wall transcripts, is still largely unknown.

Here, we reveal a novel regulatory role of Slt2p in the stabilization of cell wall transcripts under cell wall stress. Overexpression of Rbp1p leads to impaired growth of *slt2Δ* mutants and destabilization of a subset of cell wall transcripts on normal media, indicating that Rbp1p negatively regulates the stability of cell wall transcripts that are transcriptionally induced by Slt2p. Deletion of *RBP1* partially rescues the hypersensitivity of *slt2Δ* mutants and reverts the level of subset cell wall transcripts, supporting that Rbp1p acts as a negative regulator downstream of MAP kinase Slt2p signaling. We further found that Slt2p interacts with Rbp1p and enhances Rbp1p protein degradation through the C-terminal nonkinase region, which attenuates the negative control of Rbp1p on the stability of cell wall mRNAs upon cell wall stress. Given the N-terminal conserved kinase domain-regulated transcriptional activity, we reveal the posttranscriptional regulatory function of Slt2p, which is mainly exerted via its C-terminal 126 nonconserved amino acids. Slt2p-mediated attenuation of Rbp1p activity toward cell wall mRNAs could be a feedforward regulation for fine-tuning of the Slt2p MAP kinase signaling pathway. Therefore, in response to cell wall stress, Slt2p acts at both the posttranscriptional and transcriptional levels. With the simultaneous controls of distinct proteins implicated in activation of the cell wall transcriptome and in the turnover of newly synthesized cell wall mRNAs, Slt2p ensures the availability of cell wall components for further remodeling.

## Results

### RNA-binding protein Rbp1p promotes decay of a subset of cell wall transcripts

To investigate whether cell wall stress and the cell wall integrity pathway control the posttranscriptional regulatory activity of Rbp1p, wild-type and *slt2Δ* mutants were transformed individually with a high copy-based plasmid expressing the GFP-tagged *RBP1* gene under ADH promoter control [3,13,14]. Interestingly, a significant growth defect was observed for *slt2Δ* mutants expressing Rbp1p compared to wild-type cells (Figure 1A left panel). Psp1p is another negative growth regulator whose overexpression results in growth inhibition [12]. We examined whether overexpression of Psp1p can cause impaired growth of *slt2Δ* mutant cells. Intriguingly, the growth of *slt2Δ* mutants expressing Psp1p was normal compared to that of wild-type cells (Figure 1A right panel and S1A), indicating that the impaired growth phenotype of *slt2Δ* mutants caused by the expression of Rbp1p is specific and biologically significant.

**Figure 1.**
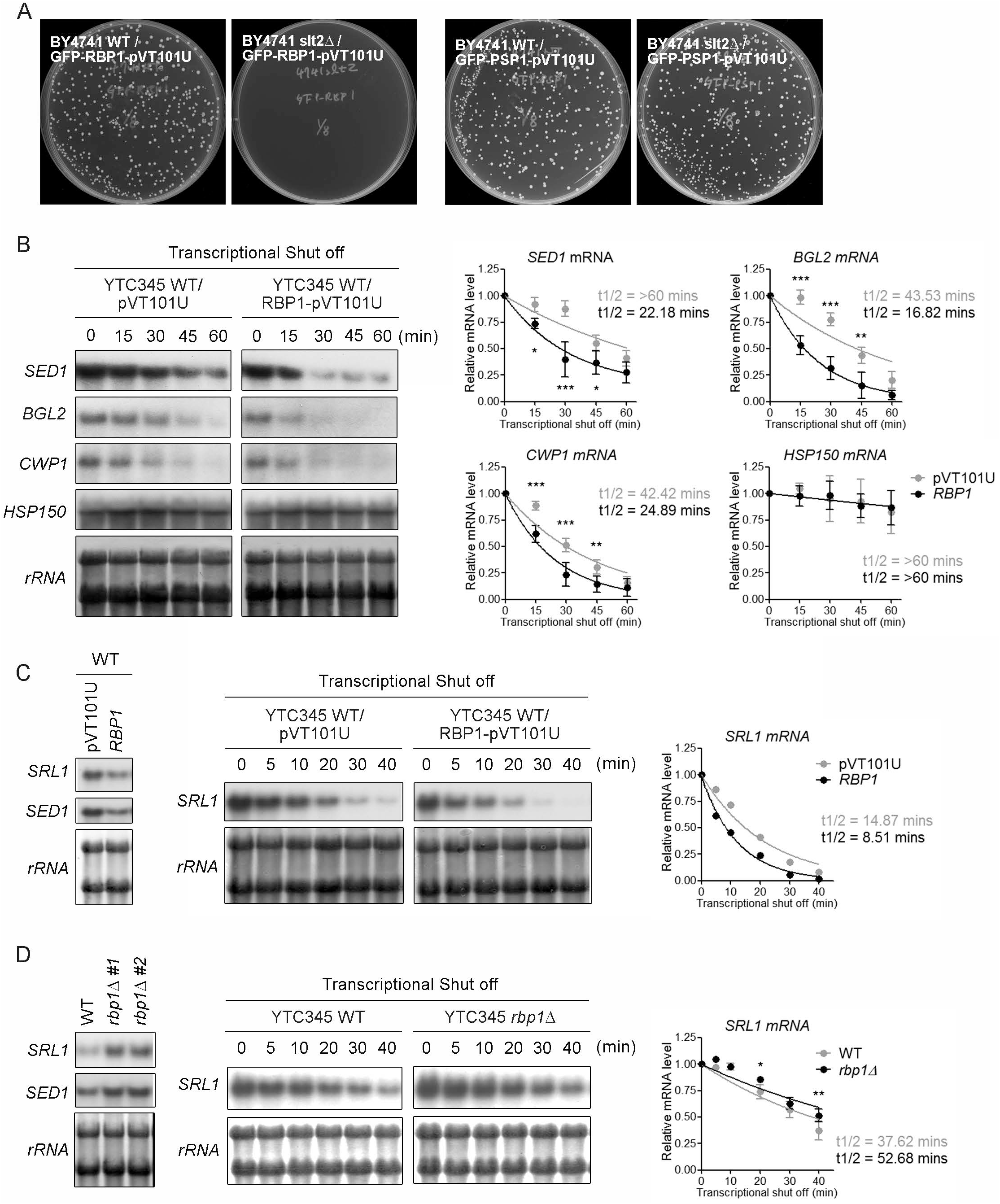
Rbp1p promotes the decay of a subset of Slt2p-regulated cell wall transcripts. (A) Overexpression of Rbp1p impaired the growth of *slt2Δ* mutant cells. BY4741 wild-type and *slt2Δ* mutant cells were transformed with GFP-RBP1-pVT101U (left panel) or GFP-PSP1-pVT101U (right panel) and spread onto SD-URA media for 2 to 3 days of culture at 30°C. (B and C) Overexpression of Rbp1p accelerates the turnover rate of a subset of cell wall mRNAs. YTC345 (*rpb1-1*) strains carrying a high-copy plasmid bearing *RBP1* or empty vector were grown in synthetic selective medium at permissive temperature to log-phase and then shifted to nonpermissive temperature to shut off transcription for the indicated times. (D) Lack of *RBP1* increased the stability of cell wall mRNA. Wild-type YTC345 (*rpb1-1*) strains or rbp1□□ mutants were grown in YPD-rich medium at permissive temperature to log-phase and then shifted to nonpermissive temperature to shut off transcription for the indicated times. Northern blotting analysis of total RNA was performed with specific probes to monitor the remaining levels of mRNA. rRNAs served as a loading control. The levels of each mRNA were quantified by ImageJ and normalized relative to those of rRNAs. t1/2 indicates the half-life of mRNAs and was calculated by one phase decay. Standard deviations are indicated. Statistical analysis using two-way ANOVA demonstrates the significance of decay kinetics.

We hypothesized that the cell growth defect caused by *RBP1* overexpression in the *slt2Δ* background might be a consequence of posttranscriptional regulation of cell wall mRNAs by Rbp1p. To examine this, the half-life of cell wall mRNAs was assessed in wild-type cells carrying the empty vector of Rbp1p. Because chemical transcriptional inhibitors are problematic for determining mRNA half-life and regulatable promoter systems are suggested [22], we took advantage of the YTC345 strain that harbors a temperature-sensitive mutant of RNA polymerase II (*rpb1-1*), allowing transcriptional shut off after shifting to the nonpermissive temperature [23]. Northern blotting analyses showed that excess Rbp1p accelerated the degradation of cell wall mRNAs, including *SED1*, *BGL2*, and *CWP1* mRNAs (Figure 1B and S1B). The turnover rates of *HSP150* mRNA, whose expression is also regulated by Slt2p during cell wall stress [20], were similar in either empty vector or Rbp1p overexpression (Figure 1B), indicating that Rbp1p may negatively regulate the stability of a subset of cell wall mRNAs.

To prove the potential role of endogenous Rbp1p in cell wall mRNA decay, we employed another cell wall mRNA, *SRL1*. We first confirmed that the steady-state level of *SRL1* mRNA was decreased upon overexpression of Rbp1p in wild-type cells (Figure 1C left panel), whereas it was increased as *SED1* mRNA in the absence of *RBP1* (Figure 1D left panel). We then determined whether the levels of *SRL1* mRNA in overexpressing or the absence of *RBP1* are attributed to Rbp1p-mediated mRNA decay. We measured the half-life of *SRL1* mRNA in the YTC345 strain in which *RBP1* was overexpressed or deleted. After shut off transcription, northern blotting analyses showed that the half-life of *SRL1* mRNA was reduced in wild-type cells overexpressing Rbp1p (Figure 1C middle and right panel) but prolonged in *rbp1Δ* mutants (Figure 1D middle and right panel). The turnover rates of *SRL1* mRNA show differences in yeast cells between cultures in synthetic and rich media (Figure 1C and 1D), which has been observed in previous reports [24,25]. Together, these results support that Rbp1p negatively regulates the stability of a subset of cell wall mRNAs.

### Deletion of RBP1 partially rescues the cell wall integrity of the slt2Δ mutant upon cell wall stress

Our above results suggest that Rbp1p may have a role in the cell wall integrity pathway and that Slt2p is required to enable yeast cells to adapt to cell wall stresses. Slt2p is the central component of the cell wall integrity response, and deletion of *SLT2* causes yeast cell hypersensitivity to cell wall antagonists, such as Congo Red, Calcolfluor White, and Caffeine [26–28]. *SED1*, *BGL2*, *CWP1*, and *SRL1*, whose mRNAs are under Rbp1p-mediated decay (Figure 1), are also known for their transcriptional upregulation by Slt2p in response to cell wall stress [21]. Considering that Rbp1p targets a subset of Slt2p-regulated cell wall mRNAs, we investigated whether loss of *RBP1* shows resistance to different cell wall stresses. We first used Congo Red, a dye that binds to chitin, since the cell wall-related transcriptional response via Slt2p has been extensively demonstrated in yeast [21]. When *rbp1Δ* mutants exert a moderate survival advantage compared to wild-type cells under Congo Red treatment (Figure 2A), deletion of *RBP1* partially suppresses the hypersensitivity of *slt2Δ* mutants to Congo Red (Figure 2B). To prevent clone biases, we analyzed two *slt2Δrbp1Δ* colonies, both of which showed similar strength of growth rescue (Figure 2B).

**Figure 2.**
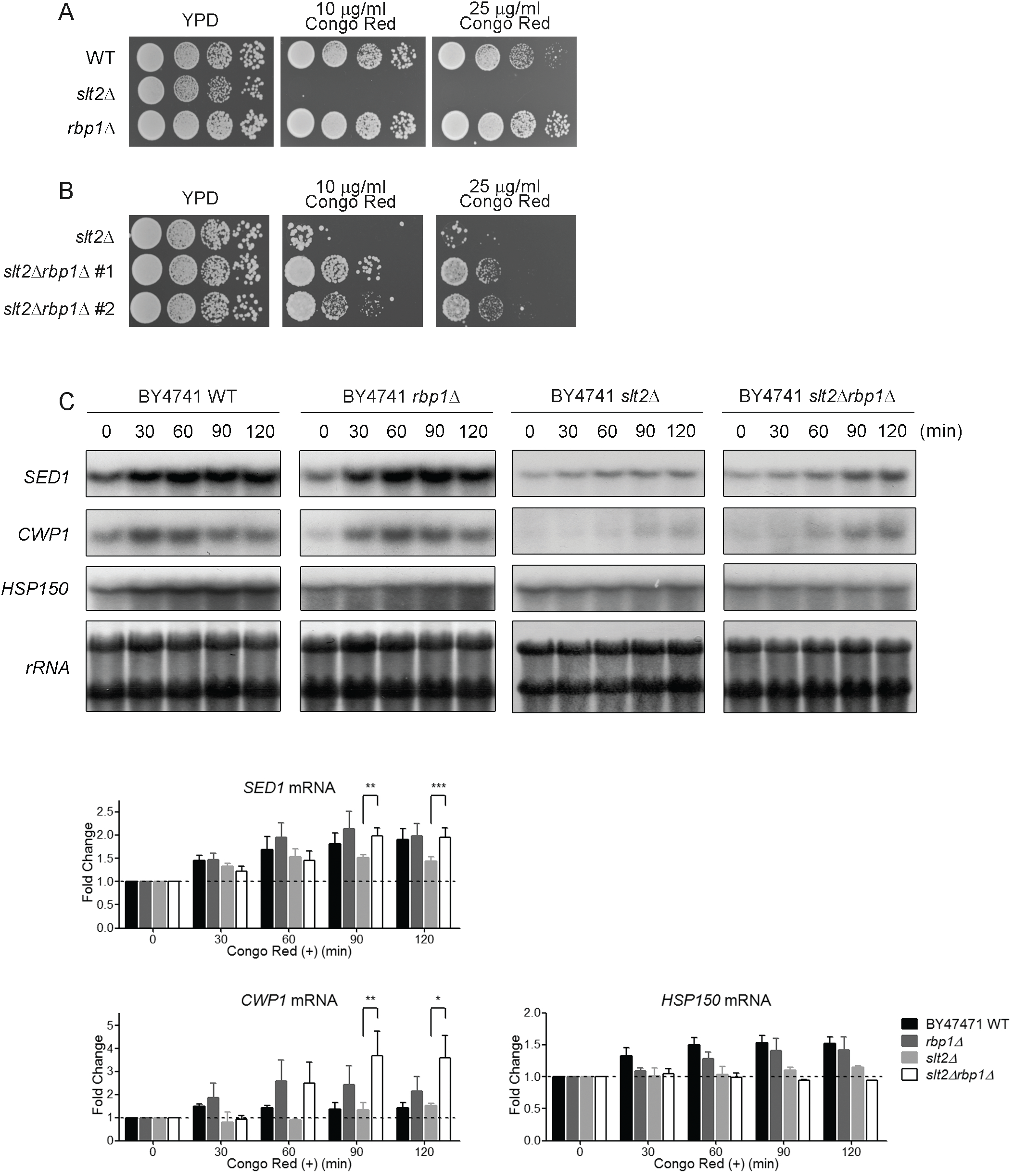
Deletion of *RBP1* partially rescues the cell wall integrity of the *slt2Δ* mutant upon cell wall stress. (A and B) Deletion of *RBP1* partially rescues the impaired growth of *slt2Δ* mutant cells. BY4741 wild-type, *slt2Δ*, *rbp1Δ*, or *slt2Δrbp1Δ* mutants were grown to log phase in rich medium, serially tenfold diluted and spotted onto rich agar medium in the presence of Congo Red (10 or 25 μg/ml) and then incubated at 30°C for 2 to 3 days. (C) Deletion of *RBP1* partially restores the induced levels of cell wall mRNAs in response to cell wall stress. BY4741 wild-type, *slt2Δ*, *rbp1Δ*, or *slt2Δrbp1Δ* mutants were grown to log phase in rich medium and then treated with 25 μg/ml Congo Red for the indicated times. Northern blotting analysis of total RNA was performed with specific probes to monitor the inducing levels of mRNA. rRNAs served as a loading control. The levels of each mRNA were quantified by ImageJ, normalized relative to those of rRNAs, and then graphed as fold change of mRNA in 0 min. Standard deviations are indicated. Statistical analysis using two-way ANOVA from two to three independent experiments demonstrates the significance of the fold change.

Caffeine is a natural purine analog. Yeast lacking *SLT2* are unable to grow on medium containing caffeine, presumably because of their weakened cell wall. However, caffeine caused cell wall perturbation largely independent of the Rlm1p transcriptional factor and excessive phosphorylation of Slt2p that is not seen in response to Congo Red [29]. When challenged with lethal doses of caffeine [29,30], *rbp1Δ* and *slt2Δrbp1Δ* mutants did not exhibit rescue of growth compared to wild-type cells and *slt2Δ* mutants, respectively (Figure S2A). Growth at elevated temperature also activates the cell wall integrity pathway [31]. Compared with wild-type cells, *rbp1Δ* mutants did not display superior growth at higher cultivation temperatures (Figure S2B). Collectively, these results indicate that Rbp1p has a stress-specific role in the Slt2p-regulated cell wall integrity response.

As stated above, Slt2p is indispensable for the upregulation of genes involved in cell wall construction and metabolism [21]. We next tested whether *rbp1*Δ suppresses *slt2*Δ lethality under Congo Red-induced cell wall stress via changes in the abundance of cell wall mRNAs. After Congo Red treatment, the transcripts of *SED1*, *CWP1*, and *HSP150* were dramatically increased in wild type and *rbp1*Δ but not in *slt2*Δ mutants (Figure 2C). Although *rbp1*Δ mutants showed a slight growth increase compared to wild-type cells in phenotypic assays (Figure 2A), the steady-state levels of cell wall mRNAs did not show a quantitative increase in *rbp1*Δ mutants compared to wild-type cells during Congo Red treatment (Figure 2C). However, in *slt2*Δ*rbp1*Δ mutants, the amounts of *SED1* and *CWP1* mRNAs were significantly increased at the late induction time points (90 and 120 min) compared with *slt2*Δ mutants, suggesting that the absence of Rbp1p partially reverts the loss of cell wall mRNAs (Figure 2C), which may correspondingly rescue the cell wall integrity of *slt2*Δ mutants upon Congo Red-induced cell wall damage (Figure 2B). Notably, *HSP150* mRNA, which is not a target of Rbp1p, did not show an increase in *slt2*Δ*rbp1*Δ mutants (Figure 2C). Together, during the Congo Red-induced cell wall integrity response, deletion of *RBP1* partially rescues the cell wall integrity of *slt2*Δ mutants, which could be attributed to increased levels of a subset of Slt2p-regulated cell wall transcripts.

### Rbp1p-targeted cell wall transcripts were stabilized upon cell wall stress

Most stress-induced transcriptionally upregulated genes have mRNA stabilization, indicating that mRNA stabilization contributes to maintaining increased mRNA levels during stress and that a common factor may exist in the coordination of both processes [16,32]. For example, Osmotic transcripts are upregulated during hyperosmotic shock by increasing the levels of transcription and mRNA half-life [16]. MAP kinase Hog1p of the HOG pathway controls both transcriptional induction and stability of osmo-responsive mRNAs [16,32]. Thus, we first investigated whether cell wall stress modulates the stability of cell wall transcripts. Wild-type YTC345 yeast cells were treated with or without Congo Red at permissive temperature for two hours and then transferred to nonpermissive temperature to shut off transcriptional activity. Northern blotting was applied to measure the half-life of cell wall mRNAs. As shown in Figure S3, cell wall transcripts, *SED1*, *CWP1*, and *BGL2* mRNAs, were stabilized after treatment with Congo Red. This result demonstrates that a subset of cell wall mRNAs was stabilized in response to cell wall stress.

We next examined whether MAP kinase Slt2p is involved in stress-induced stabilization of cell wall transcripts. To test this hypothesis, we compared the turnover rates of those mRNAs that responded quickly to Congo Red treatment in YTC345 wild-type cells and *slt2*Δ mutants. As shown in Figure 3A, upon cell wall stress, the average half-life of *SED1* mRNA was more than 60 minutes in wild-type cells, while it was reduced to approximately 30 minutes in *slt2*Δ mutants. A comparable half-life reduction was observed with *BGL2*, *CWP1*, and *HSP150* mRNAs, agreeing that MAP kinase Slt2p regulates cell wall gene expression at both transcriptional and transcript turnover levels.

**Figure 3.**
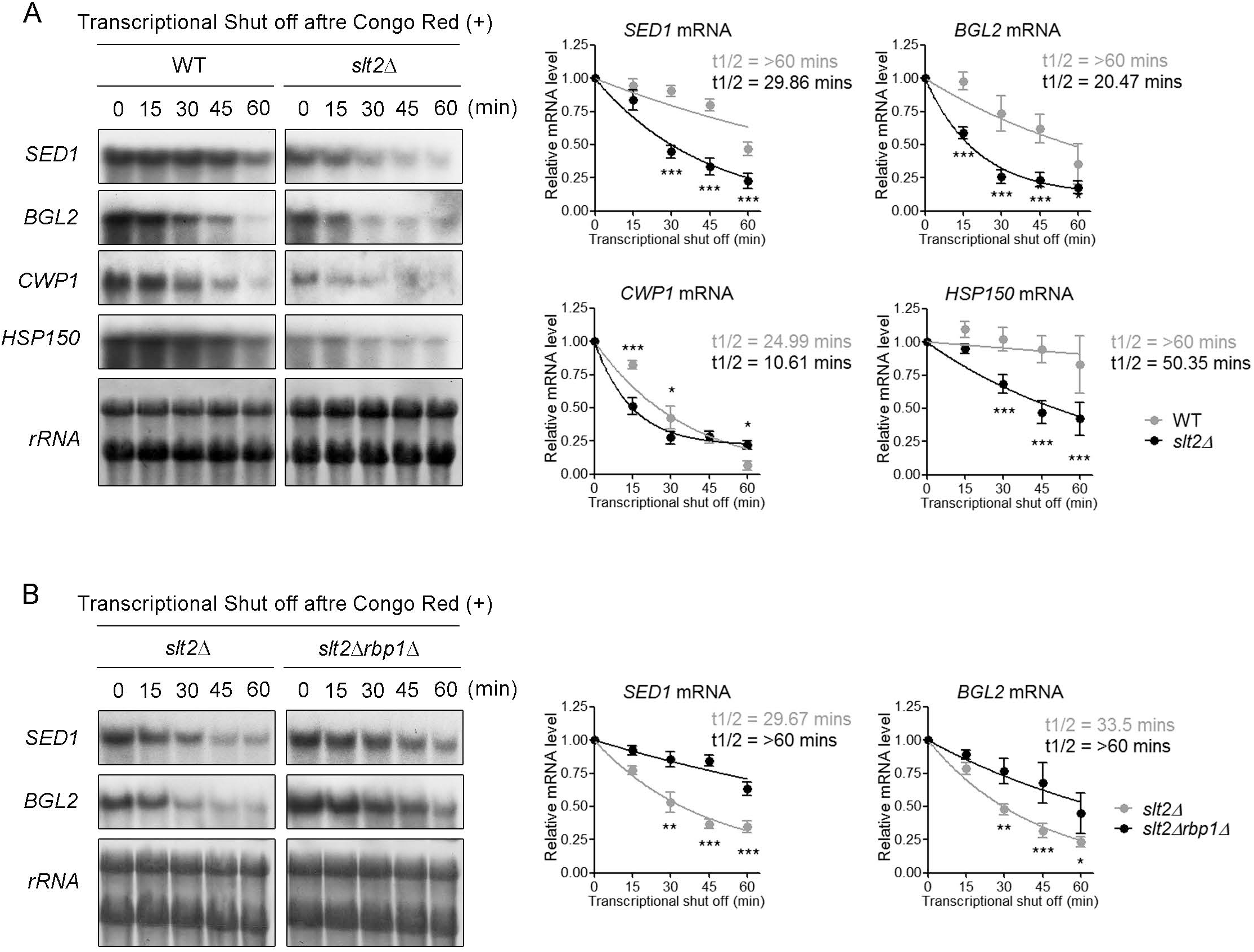
Stabilization of a subset of cell wall transcripts in response to cell wall stress depends on Slt2p. (A) Slt2p is required for stabilization of cell wall mRNAs during cell wall stress. YTC345 (*rpb1-1*) wild-type or *slt2Δ* mutants were grown in rich medium to log-phase and treated with 25 μg/ml Congo Red for 2 hr. (B) Deletion of *RBP1* rescues the instability of cell wall mRNAs caused by the absence of Slt2p. YTC345 (*rpb1-1*) *slt2Δ* or *slt2Δ*rbp1*Δ* mutants were grown in rich medium to log-phase and treated with 25 μg/ml Congo Red for 2 hr. After treatment, the cells were shifted to a nonpermissive temperature to shut off transcription for the indicated times. Northern blot analysis, quantification and statistical analysis of (A) and (B) were performed as described in Figure 1.

Removing *RBP1* relieved the loss of cell wall mRNAs in *slt2*Δ mutants treated with Congo Red (Figure 2C). Similar suppression activity of *rbp1*Δ mutants was observed when examining the half-life of Rbp1p-targeted cell wall mRNAs in *slt2Δrbp1*Δ mutants. Upon Congo Red treatment, the half-life of *SED1 and BGL2* mRNAs in *slt2Δrbp1*Δ mutants increased to approximately 60 minutes, similar to that in wild-type cells, while only approximately 30 minutes in *slt2*Δ mutants (Figure 3B). We also compared the stability of cell wall transcripts in both wild-type and *rbp1*Δ mutant cells upon cell wall stress. However, the half-life of the examined cell wall mRNAs was similar between wild-type and *rbp1*Δ mutant cells (data not shown), reminiscing that in Figure 2C, the amount of cell wall mRNAs did not show a significant increase in *rbp1*Δ mutants compared to wild-type cells during Congo Red treatment. We further discuss it in the Discussion. Altogether, the reduced cell wall mRNA stability in *slt2*Δ mutants reverted in *slt2*Δ*rbp1*Δ mutants indicated that Rbp1p-mediated cell wall mRNA decay is critical for the regulation of gene expression during cell wall stress and suggested that Rbp1p plays a role downstream of Slt2p in regulating the stabilization of cell wall mRNAs.

### Slt2p directly interacts with Rbp1p in response to cell wall stress

Slt2p contains a highly conserved N-terminal MAP kinase domain, which shares a similar activation mechanism from yeast to humans. However, the extended C-terminal region of Slt2p is functionally unclear but is assumed to regulate its own kinase activity [33]. As the genetic interaction of Rbp1p and Slt2p during cell wall stress is recognized, we further investigated whether Rbp1p is a potential downstream effector of Slt2p in response to cell wall stress. We first observed that full-length Slt2p showed a strong interaction with Rbp1p in a yeast two-hybrid assay (Figure 4A and 4B). Deleting the last 126 amino acids of Slt2p (Slt2p-dC126) completely abolished the Slt2p-Rbp1p interaction, while Slt2p with a slightly shorter deletion (Slt2p-dC115) showed no defect in binding Rbp1p (Figure 4A and 4B). However, expression of the C-terminal 126-residue of Slt2p alone is not sufficient for its interaction with Rbp1p (Figure S4), implying that the binding of Slt2p to Rbp1p requires the Slt2p kinase domain and is stabilized by the accessory of the Slt2p C-terminus.

**Figure 4.**
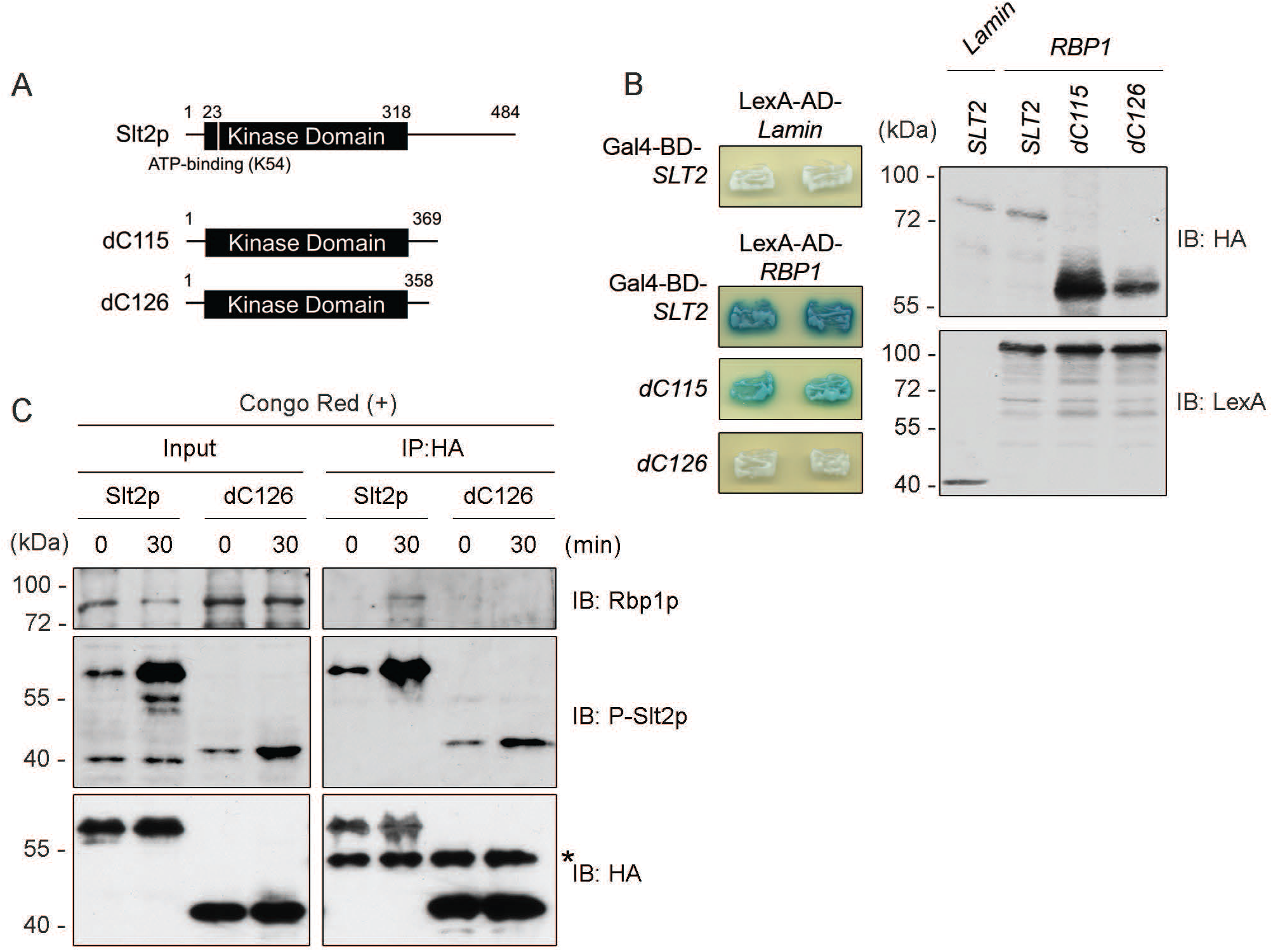
Slt2p interacts with Rbp1p in response to cell wall stress. (A) Schematic representation of the Slt2p protein domain structure and C-terminal truncated variants. (B) Slt2p interaction with Rbp1p requires its C-terminal 126 amino acids in a yeast two-hybrid assay. YEM1α cells carrying LexA- and Gal4AD-based fusion constructs as indicated were assayed for β-galactosidase activity. Western blotting was used to analyze the expression levels of the indicated fusion proteins. (C) Slt2p interacts with Rbp1p endogenously in response to cell wall stress through the C-terminus. BY4741 wild-type cells chromosomally expressing Slt2p-3HA or Slt2p-dC126-3HA were treated with 25 μg/ml Congo Red for the indicated times. Cell extracts were used for immunoprecipitation with anti-HA antibody-conjugated beads, followed by western blotting analysis with the indicated antibodies.

Next, we tested whether Slt2p interacts with Rbp1p in a C-terminal-dependent manner in vivo by coimmunoprecipitation experiments in the yeast strain (Slt2p-HA) with 3xHA tagged into the chromosomal locus of *SLT2*. As expected, Rbp1p was discovered in Slt2p-HA immunoprecipitates 30 minutes after Congo Red treatment, indicative of an endogenous interaction of Slt2p and Rbp1p (Figure 4C). Notably, no interaction of Rbp1p and Slt2p was observed under nonstressed conditions (Congo Read treatment, 0 min), suggesting that this interaction is accompanied by the activation of cell wall stress signaling, reflected by the phosphorylation status of Slt2p (Figure 4C). Activated Slt2p was detected with antibodies directed against mammalian phosphorylated ERK1/2 [27]. Consistent with the results from yeast two-hybrid assays, Slt2p with a deleted C-terminus (Slt2p-dC126-HA) was deficient in the endogenous interaction between Slt2p and Rbp1p upon Congo Red treatment (Figure 4C). Together, our results showed that Rbp1p is a novel Slt2p-interacting protein under the stimulation of cell wall stress and that the C-terminal region of Slt2p is essential for this interaction.

### Slt2p interaction mediates cell wall stress-induced degradation of Rbp1p

Considering that Rbp1p has an opposing effect to Slt2p on cell wall mRNAs, we hypothesized that the mRNA decay activity of Rbp1p was attenuated upon cell wall stress. We examined the protein level of Rbp1p under normal and cell stress conditions. Rbp1p protein gradually decreased after treatment with Congo Red, while it showed a comparable amount under normal conditions (Figure S5A), indicating that cell wall stress induces a decrease in Rbp1p protein. With no fluctuation of *RBP1* transcripts observed during the course of Congo Red treatment (Figure S5B), we further examined whether Rbp1p was targeted for degradation during cell wall stress. Normal growth or Congo Red-treated wild-type cells were subjected to cyclohexmide chase experiments, and the half-life of endogenous Rbp1p was determined accordingly (Figure 5A). The half-life of Rbp1p protein in normal growth is approximately 16 minutes, while the half-life of Rbp1p protein in yeast exposed to Congo Red is reduced to approximately 9 minutes, demonstrating that Rbp1p was subjected to protein degradation in response to cell wall stress (Figure 5A). These results suggest that cell wall stress could attenuate the RNA decay activity of Rbp1p through protein degradation.

**Figure 5.**
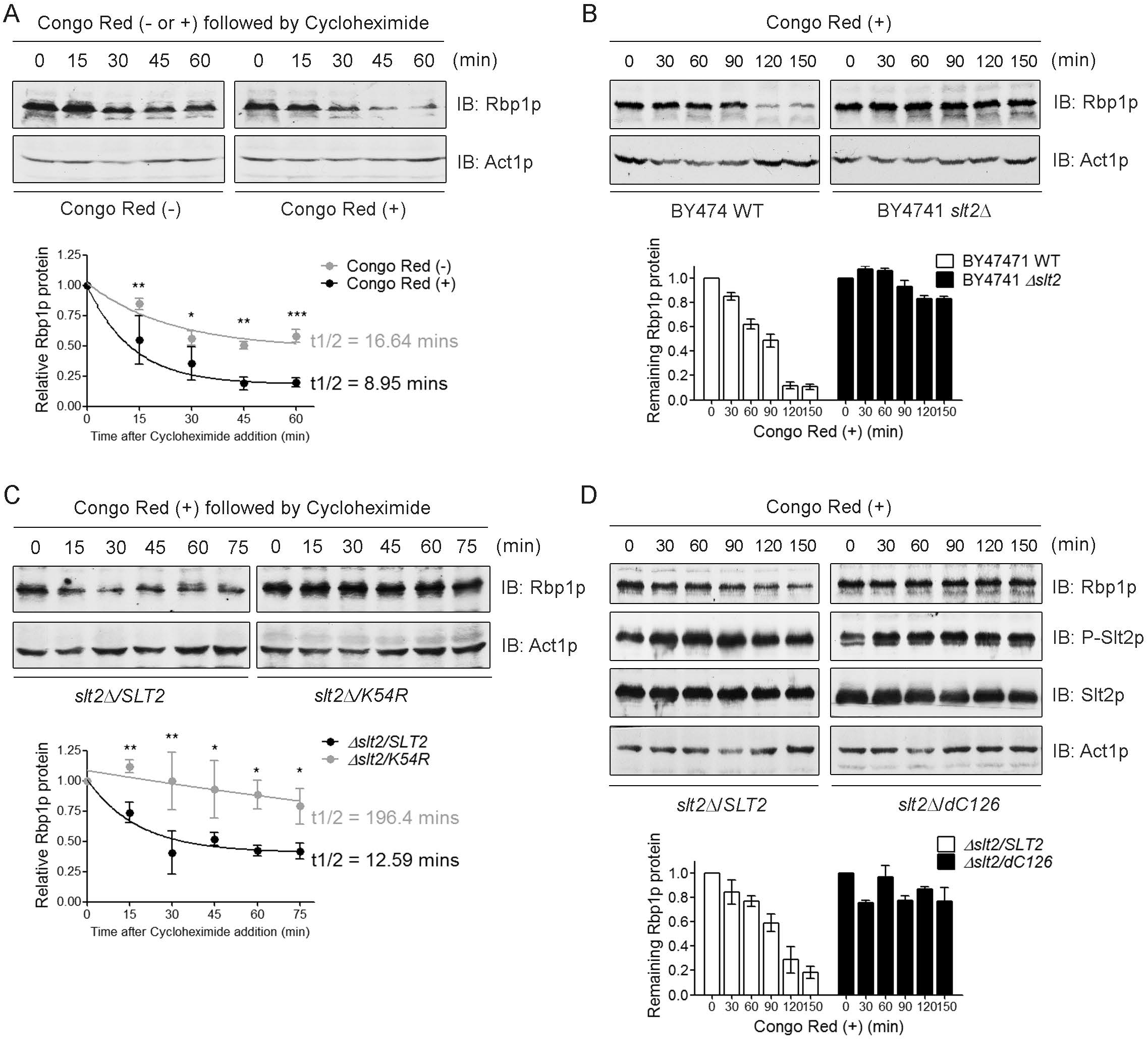
Slt2p mediates the degradation of Rbp1p in response to cell wall stress. (A) The turnover rate of Rbp1p is promoted by cell wall stress. Wild-type cells were treated with either Congo Red or not for 2 hr and then incubated with 100 μg/ml cycloheximide for the indicated times. (B) Rbp1p decreased significantly 120 minutes after Congo Red treatment in wild cells but remained unchanged in *slt2*Δ mutants. (C) The degradation of Rbp1p in response to the cell wall required Slt2p kinase activity. *slt2*Δ mutants expressing wild-type *SLT2* or the kinase-dead *K54R* mutant were pretreated with 25 μg/ml Congo Red for 2 hrs, followed by incubation with 100 μg/ml cycloheximide for the indicated times. (D) Slt2p-dC126 fails to decrease Rbp1p protein during cell wall stress. *slt2*Δ mutants expressing Slt2p or Slt2p-dC126 were treated with Congo Red for the indicated times. Western blots show Rbp1p protein levels in BY4741 cells. The levels of Rbp1p were quantified by ImageJ and normalized relative to those of Act1p, and then graphed as relative levels of Rbp1p in 0 min (B, D) or t1/2 indicated the half-life of Rbp1p and was calculated by one phase decay (A, C). Standard deviations from three independent experiments are indicated. Statistical analysis using two-way ANOVA demonstrates the significance of decay kinetics.

The Slt2p MAP kinase cascade has been shown to mediate the destruction of the downstream transcriptional repressor C-type cyclin in response to oxidative stress (Krasley et al. 2006). Next, we investigated whether Slt2p is required for the reduction of Rbp1p protein in yeast cells subjected to cell wall stress. The levels of Rbp1p protein were reduced in wild-type yeast cells following treatment with Congo Red, while Rbp1p protein levels remained unchanged in *slt2*Δ mutant cells even two hours after Congo Red treatment (Figure 4B). This finding suggests that Slt2p and the MAP kinase cascade may transduce the cell wall stress degradation signal for Rbp1p.

To obtain better insight into the mechanism underlying this process, we next asked whether the kinase activity of Slt2p is involved in Rbp1p degradation in response to cell wall stress. Compared with the expression of wild-type Slt2p, the expression of the Slt2p kinase-dead version (Slt2p-K54R) in *slt2*Δ mutant cells neither rescued the cell growth defects properly (Figure S6A and S6B) nor promoted Rbp1p reduction and degradation efficiently after Congo Red treatment (Figure 4C and S4C). Detection of phosphorylation at Thr190 and Tyr192 in Slt2p-K54R cells (Figure S4C) [34] indicated that MAP kinase cascade signaling is intact and that the kinase activity of Slt2p is indispensable for Rbp1p degradation.

Similar defects in cell growth and Rbp1p degradation are exhibited in *slt2*Δ mutant cells expressing inactive Slt2p-TA/YF (Figure S6A, S6B, and S6D), whose activation loop phosphoracceptors, Thr190 and Tyr192, are mutated to avoid receiving signals from upstream MAPKKs Mkk1/Mkk2 (Figure S6A) [34]. Failure to detect phospho-Slt2p along with stabilization of Rbp1 protein after Congo Red treatment highlights the importance of MAP kinase cascade signaling in Rbp1p degradation (Figure S6D). Given that the kinase activity of Slt2p is essential for the activation of the downstream transcription factor Rlm1p under cell wall stress [35–37], here, we demonstrated that the cell wall integrity signaling and kinase activity of Slt2p are both required for the degradation of the Rbp1p protein in response to cell wall stress.

To further determine whether the Slt2p-Rbp1p interaction mediated this degradation process, we examined endogenous Rbp1p protein levels in *slt2*Δ mutant cells expressing wild-type Slt2p or truncated Slt2p-dC126 upon Congo Red treatment. Western blot analysis showed that Rbp1p protein from cells expressing Slt2p-dC126 was more stable than that from cells expressing Slt2p after Congo Red treatment for more than 2 hours (Figure 5D). Interestingly, Slt2p-dC126 is properly phosphorylated in response to cell wall stress signals, an event that reflects the activation of Slt2p, indicating that the removal of the C-terminal Rbp1p-interacting region of Slt2p has no effect on biochemical behavior in response to cell wall stress. Together, our observations suggested that Slt2p downregulates the protein level of Rbp1p upon cell wall stress and that the interaction between Slt2p and Rbp1p is indispensable for this process.

### Slt2p stabilizes cell wall transcripts upon cell wall stress via interaction with Rbp1p

To prove that MAP kinase Slt2p stabilizes a subset of cell wall transcripts during cell wall stress by downregulating Rbp1p activity, we first examined the transcriptional activity of Slt2p-dC126 upon the Congo Red-induced cell wall integrity response. The levels of *SED1*, *BGL2*, *CWP1* and *HSP150* mRNAs in *slt2Δ* mutant cells expressing wild-type Slt2p were fairly induced after Congo Red treatment for 2 hours (Figure 6A). However, in *slt2Δ* mutant cells expressing Slt2p-dC126, the mRNA levels of *SED1* and *BGL2* were induced but much less than those in cells expressing Stl2p, especially after treatment with Congo Red for 90 and 120 mins (Figure 6A). The reduced steady-state levels of a subset of cell wall mRNAs at late time points of cell wall stress leads us to speculate that this could be caused by the loss of interaction of Slt2p-dC126 with Rbp1p to attenuate Rbp1p-mediated cell wall mRNA decay activity.

**Figure 6.**
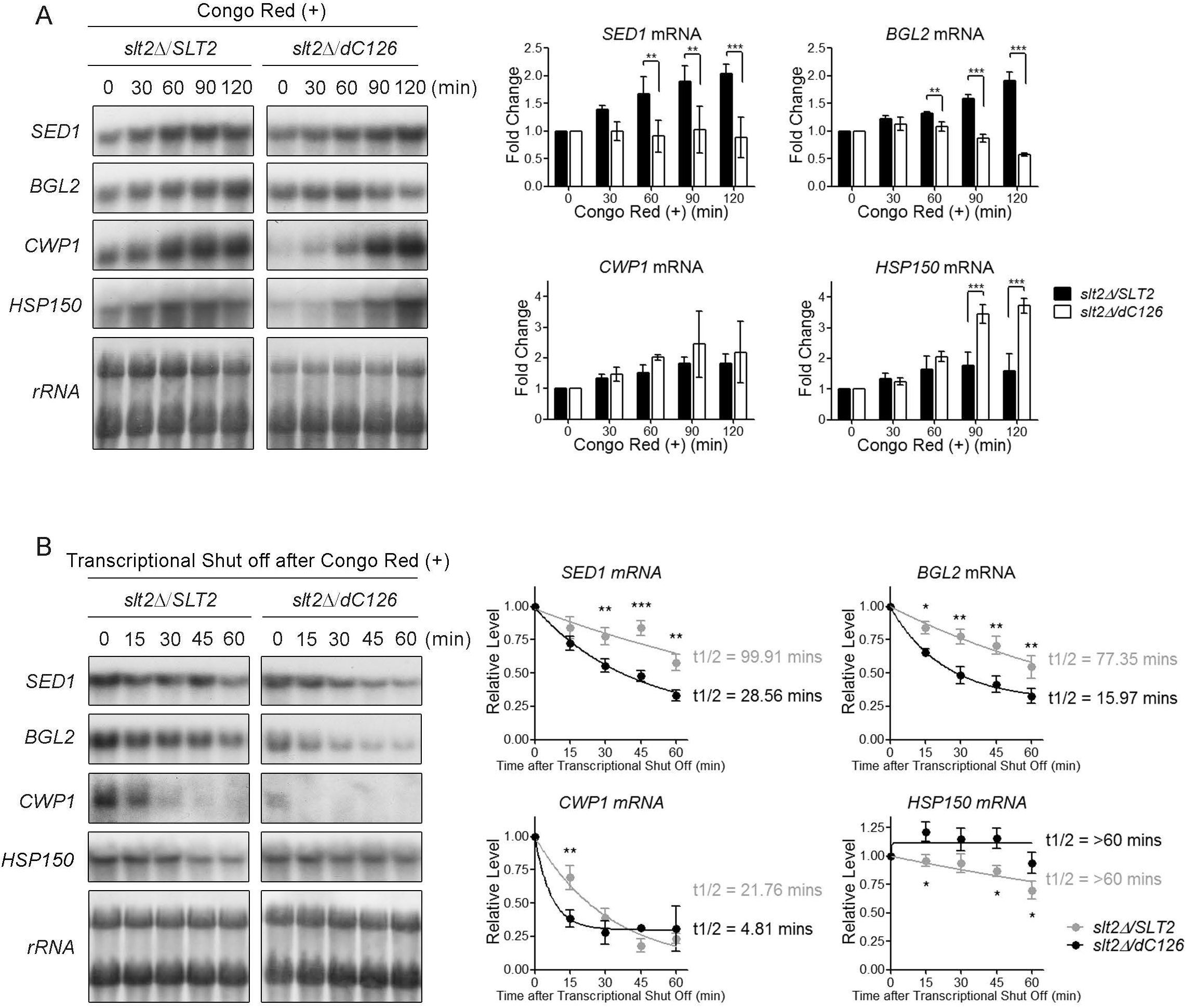
The C-terminal Rbp1p-interacting region of Slt2p functions in maintaining the stabilization of Rbp1p-targeted cell wall mRNAs. (A) Slt2p lacking C-terminal 126 amino acids failed to fully restore the induced levels of Rbp1p-targeted cell wall mRNAs in response to cell wall stress. BY4741 sl*t2Δ* mutants expressing full-length *SLT2* or *SLT2-dC126* were grown to log phase in synthetic selection medium and then treated with 25 μg/ml Congo Red for the indicated times. (B) C-terminal 126 amino acids of Slt2p function in maintaining the stability of cell wall mRNAs. YTC345 (*rpb1-1*) *slt2Δ* mutants expressing full-length *SLT2* or *SLT2-dC126* were grown in synthetic selection medium at permissive temperature to log-phase, treated with 25 μg/ml Congo Red for 2 hrs and then shifted to nonpermissive temperature to shut off transcription for the indicated times. Northern blotting analysis of total RNA was performed with specific probes to monitor the inducing levels of mRNA. rRNAs served as a loading control. The levels of each mRNA were quantified by ImageJ and normalized relative to those of rRNAs, and then graphed as fold change of mRNA in 0 min (A) or t1/2 indicated the half-life of mRNAs and was calculated by one phase decay (B). Standard deviations are indicated. Statistical analysis using two-way ANOVA from two to three independent experiments demonstrates the significance of fold change (A) and decay kinetics (B).

To test this possibility, we examined the turnover rates of cell wall mRNAs in YTC345*slt2Δ* mutants expressing full-length Slt2p or Slt2p-dC126 upon cell wall stress. We speculated that Slt2p lacking the C-terminal Rbp1p-interacting region would not restore the stabilization of cell wall mRNAs as efficiently as the wild-type protein because of its failure to interact with endogenous Rbp1p (Figure 3C) to mediate Rbp1p protein degradation (Figure 5D). Northern blotting showed that in YTC345*slt2Δ* mutants expressing wild-type Slt2p, *SED1* mRNA was stable, and the turnover rate was restored to more than 60 min upon cell wall stress (Figure 6B). However, in cells expressing truncated Slt2p-dC126, *SED1* mRNA was more liable and reduced its turnover rate to lower than half after Congo Red treatment (Figure 6B). The same conclusion comes with *BGL2* and *CWP1* mRNAs (Figure 6B). These results demonstrated that 126 C-terminal amino acids, Rbp1p-interacting regions, are required for Slt2p to stabilize a subset of stress-induced cell wall mRNAs.

Interestingly, in contrast to Rbp1p-targeted cell wall transcripts, the amount and half-life of *HSP150* mRNA increased in *slt2Δ* mutants expressing Slt2p-dC126 compared to those expressing full-length Slt2p (Figure 6A and 6B), implying that for different subsets of cell wall transcripts, Slt2p may exert different regulatory mechanisms via its C-terminal extension.

We further examined whether the C-terminal deleted variants of Slt2p still possess biological functions. Phenotypical rescue experiments showed that Slt2p-dC126, as Slt2p, fully complemented the death phenotype of *slt2Δ* mutant cells to grow in the presence of Congo Red (Figure S7B). In contrast, expression of Slt2p-dC142 cannot rescue the growth defeat of *slt2Δ* mutant cells (Figure S7B), regardless of equal protein expression levels (Figure S7A). Interestingly, compared to caffeine treatment (Figure S7C), Slt2p-dC126 shows slightly better growth rescue activity upon Congo Red-induced cell wall damage (Figure S7B and S7C), implying that the C-terminal 126 residues of Slt2p play different regulatory activities when facing different environmental conditions.

## Discussion

Genes that are transcriptionally upregulated by stresses usually have better mRNA stabilities, implying the need for common factors that coordinate both processes [16,32]. Microarray approaches have shown that individual RNA-binding proteins specifically associate with a subset of mRNAs that are functionally related. Hence, an RNA-binding protein may coordinate the stability of a group of mRNAs rather than targeting an individual mRNA [38–40]. These observations favor the scenario that functionally related genes are coregulated as a posttranscriptional operon or RNA regulon by specific RNA-binding proteins during stresses [41,42]. Here, we provide an example that meticulous and organized control of mRNA abundance is critical for yeast cells to adapt to new environmental conditions. In addition to alterations in cell wall gene transcription, changes in the cell wall mRNA stability of particular subsets also contribute to cell wall integrity responses. Taking advantage of blocking RNA polymerase II, we showed that changes in the stability of cell wall mRNAs are coupled with their transcription rates during Congo Red-induced cell wall stress.

We have previously shown that Rbp1p binds to the 3’-UTR of mitochondrial *porin* mRNA and then promotes the turnover of *porin* mRNA [13]. Rbp1p-mediated *porin* mRNA decay requires the direct interaction of Rbp1p with Dhh1p, a decapping activator and RNA helicase [3], which subsequently recruits the Xrn1p-dependent decay machinery [14]. Interestingly, our preliminary works show that Rbp1p may regulate the stability of several cell wall mRNAs in cooperation with two decapping activators, Pat1p and Lsm1p, but not Dhh1p, suggesting two classes of Rbp1p-mediated mRNA decay processes by partnering with different decapping activators. Lack of Pat1p abolished the formation of P-bodies during Congo Red-induced cell wall stress [17]. Rbp1p translocates to P-bodies under glucose deprivation or KCl treatment, and the failure of P-body movement disrupts the regulation of porin mRNA by Rbp1p [14]. Whether Slt2p has an additional function in Rbp1p localization is still an open question.

In eukaryotes, mRNAs that encode proteins belonging to the same biological process usually have coordinated decay as well [43,44]. It is also known that RNA-binding proteins tend to bind mRNAs sharing similar cellular functions [39,40]. The specificity of selective mRNAs for degradation is recognized via the cis-acting regulatory sequences residing in the mRNA untranslated regions [45]. To better understand the mechanism of Rbp1p-mediated mRNA degradation, it would be noteworthy to determine if the Rbp1p-targeting cell wall mRNAs harbor any common sequence motifs by which their stabilities are modulated. In addition, the level of a transcript within a cell depends on its rate of synthesis and rate of decay. Previous reports provide evidence that these two processes are integrated, and transcription factors and promoter elements of genes can directly influence the relative stability of transcripts that they induce [46,47]. Whether upstream activating sequences (UAS) or promoter elements of Rbp1p-targeted cell wall transcripts display conservation and Rbp1p-binding accessibility is also a tempting question for further investigation.

Erk1/2 and Erk5 are two human orthologs of yeast Slt2p. Sequence alignment of yeast Slt2p and human Erk1/2 and Erk5 represents a large sequence extension from the C-terminus of the MAP kinase domain of Slt2p and Erk5 but not Erk1/2. Expression of human Erk5 in yeast complements the lack of Slt2p and confers Slt2p function, including temperature and caffeine sensitivity [48,49]. Human Erk1/2 lacking a longer C-terminal tail was spontaneously phosphorylated when expressed in yeast and partially rescued the cell wall damage in *slt2Δ* mutants treated with caffeine [33].

Removal of the C-terminal 126 residue of Slt2p makes Slt2p-dC126 mimic Erk2 and spontaneously phosphorylates and partially rescues phenotypic activity during caffeine challenge [33,34]. In agreement with previous studies, our studies confirmed that Slt2p-dC126 shows spontaneous phosphorylation and renders cell integrity activity under Congo Red treatment (Figure 5D, S7A and S7B). The Slt2p MAP kinase cascade has been shown to mediate destruction of a transcriptional repressor C-type cyclin in response to oxidative stress [50]. Here, we demonstrated that Slt2p downregulates the protein level of the RNA-binding protein Rbp1p upon cell wall stress to inactivate the RNA decay activity of Rbp1p. We found that under normal growth conditions, a lack of *RBP1* leads to stabilization of the cell wall transcript (Figure 1D). However, upon cell wall stress, Rbp1p protein undergoes degradation, and the amount (Figure 2C) and half-life (data not shown) of cell wall transcripts did not significantly increase in *rbp1Δ* mutant cells compared to wild-type cells. It could be explained that Slt2p-mediated Rbp1p degradation in wild-type cells is as efficient as manual deletion of *RBP1* to cause an effect on the stability of cell wall transcripts, therefore leading to no difference in RNA level and half-life observation.

Several mammalian signal transduction pathways, including p38 MAPK/SPAK, phosphoinositide-3-kinase (PI3K) and mammalian target of rapamycin (mTOR), are known to regulate RNA stability and decay; however, studies addressing the connections between these signaling pathways and RNA-binding proteins are limited [51,52]. For example, p38 MAP kinase is an inflammatory response factor that regulates the stability of many inflammatory mRNAs. MK2, a downstream kinase effector of p38, directly phosphorylates the mRNA-destabilizing protein tristetraprolin (TTP). The phosphorylated TTP has changes in stability and subcellular localization, which in turn reduces the destabilizing activity of TTP in ARE-containing cytokine mRNAs [53,54]. Yeast MAP kinase Slt2p is known for its role in modulating transcription programs to overcome the duration of cell wall stresses. Slt2p controls several transcription factors (e.g., Rlm1p and SBF complex Swi4/Swi6p) both catalytically and noncatalytically, which in turn activate the transcription of cell wall genes [18,55]. Our results demonstrate that Slt2p has combined effects on cell wall genes: inducing their transcription and preventing the gene products from degradation posttranscriptionally. The latter requires Slt2p kinase activity to mediate the degradation of Rbp1p. Using the Eukaryotic Linear Motif (ELM) resource, we mapped and narrowed out three putative Slt2p-mediated phosphorylation sites on Rbp1p. Rbp1p could be a new substrate of Slt2p kinase, and the Slt2p-dependent phosphorylation of Rbp1p may be crucial for its stability during cell wall stress. Alternatively, Slt2p-dependent phosphorylation of Rbp1p may change its affinity for mRNAs or its protein-protein interactions with mRNA decay components. Altogether, stress-induced cell wall mRNAs are preserved for further translation to supplement the damaged cell wall.

Slt2p may regulate mRNAs at the posttranscriptional level via mechanisms other than modulating mRNA stability. It has been reported that Slt2p phosphorylates the RNA-binding protein Nab2p following heat shock stress. Nab2p, as a poly(A)mRNA-binding protein, subsequently dissociates from the mRNA export receptor Mex67p, which increases the nuclear retention of poly(A)mRNAs but favors the export of heat shock mRNAs necessary for thermotolerance [56]. Nevertheless, here, we have proposed a novel regulatory process of the cell wall stress response via the Slt2p-Rbp1p interaction; to better understand the mechanistic details requires further study.

## Materials and Methods

### Strain, media, and plasmid construction

The yeast strains used in this study are listed in Supplementary Table I. Yeast cells were grown either in rich medium containing 1% yeast extract, 2% peptone and 2% glucose or in synthetic media containing 0.67% yeast nitrogen base (without amino acids) and 2% glucose supplemented with the appropriate nutrients. Yeasts were transformed by the lithium acetate method [57]. The *SLT2* gene was disrupted in YTC345 using a Kan disruption cassette amplified by PCR from pFA6-kanMX6 [58]. Strains expressing Slt2p-3HA or Slt2p-dC126-3HA were obtained through insertion of a 3HA-HIS cassette amplified from pFA6a-3HA-His3MX6 [58]. Disruption or insertion of each cassette was verified by western blotting. Plasmids were constructed and are listed in Supplementary Table II.

### Phenotype Analysis

Yeast cultures were grown in YPD-rich or synthetic selection medium to mid-log phase (OD600 of ~1.0). Serial 10-fold dilutions were prepared. Five microliters from each dilution was spotted on YPD plates supplemented with Congo Red, caffeine or synthetic medium plates containing 2% glucose as the carbon source, incubated at 30°C for days, and photographed.

### Cycloheximide Chase Assay

Growth yeast cells in YPD-rich medium or synthetic selection medium in flasks at 30° C until the cell density reached OD600 ~1.0. After cultivation, yeast cells were treated with Congo Red at a working concentration of 25 μg/ml for 2 hours, followed by the addition of cycloheximide to 100 μg/ml. Immediately after adding cycloheximide and equilibrating cell suspensions for 5 min at 30°C, 1 ml of yeast cell suspension was harvested every 15 minutes with added cycloheximide to the microcentrifuge tube, centrifuged to pellet down yeasts, and frozen to −20°C. Yeast proteins were extracted using glass beads and TCA methods.

### Yeast two-hybrid assay

The yeast strain YEM1a was cotransformed with different combinations of bait (pEG202) and prey (pJG4-5) plasmids, and b-galactosidase plate assays were performed by streaking transformants onto SC-Trp-His plates containing 2% galactose and 80 mg/ml X-Gal (5-bromo-4chloro-3-indolyl-b-D-galactoside). The plates were then incubated at 30°C for 2–3 days.

### Yeast cell extract preparation and western blotting

Extracts were obtained from ~3 OD600 of yeast cells, suspended in 5% TCA and processed by vigorous vortexing with glass beads. Cell debris was collected by centrifugation at 13 000 rpm for 10 min, washed with water to remove residual TCA, centrifuged at 13 000 rpm for 10 min, suspended in SDS-loading buffer and then heated at 95°C for 5–10 min. For western blotting, all cell extracts were run on 9% SDS–polyacrylamide gels. Proteins were then transferred to nitrocellulose membranes and probed with the indicated antibodies. Act1p was used as a loading control.

### Northern blotting and mRNA decay assay

For steady-state mRNA analysis, cells were grown in synthetic medium lacking the indicated nutrients and containing 2% glucose to log phase. For mRNA decay analysis, the yeast strain YTC345 carrying a temperature-sensitive RNA polymerase II allele (rpb1-1) was grown at 25°C in synthetic medium lacking the indicated nutrients and containing 2% glucose until an OD600 of ~1.25 was attained and then shifted to a 37°C water bath shaker to block transcription activity of RNA polymerase II. Aliquots were collected at the indicated time points after transcription shut-off for total RNA isolation and northern blot analysis. Total RNA was prepared by the hot acid phenol method, and 10 mg of each total RNA sample was separated by 1.2% agarose gel electrophoresis in the presence of 3.7% formaldehyde. Transfer to nylon membrane (Millipore) was achieved by capillary action with 20X SSC. Blots were probed with 32P-radiolabeled riboprobes directed against the genes as indicated. The level of mRNA in the northern blots was determined by quantifying the intensity of bands using ImageJ software in three independent experiments, normalized against the intensity of rRNA, and graphed with Microsoft Excel.

### Immunoprecipitation

Exponentially growing cells (OD600 ~10) were disrupted with glass beads in 0.4 ml of extraction buffer [25 mM HEPES–KOH, pH 7.5, 75 mM KCl, 2 mM MgCl2, 0.1% NP-40, 1 mM DTT, 0.2 mg/ml heparin, 20 U/ml DNase (TaKaRa) and 10 mg/ml aprotinin, leupeptin, and pepstatin]. Extracts were cleared by centrifugation at 4000 g for 10 min. Monoclonal anti-HA antibody-conjugated agarose beads (mouse monoclonal anti-HA-agarose antibody) (Sigma #A2095) were added to the cleared extracts and incubated at 4°C for 4 h. Beads were washed four times with wash buffer (25 mM HEPES-KOH, pH 7.5, 75 mM KCl, 2 mM MgCl2, 0.1% NP-40), and the bound complexes were eluted with 50 mM Tris-HCl, pH 8.0, 100 mM NaCl, 10 mM EDTA, and 1% SDS for 10 min at 65°C. HA-tagged proteins from cell extract and immunoprecipitate were separated on a 9% SDS–PAGE gel, blotted and hybridized with anti-Rbp1p or anti-Slt2p antibody for the presence of proteins.

### Statistical Analysis

GraphPad Prism 5 was used to analyze the significance of repeated experiments.

## Acknowledgments

The authors thank Drs. Woan-Yuh Tarn and Tien-Hsien Chang for critical reading of the manuscript.

## Funding

Funding for open access charge: National Health Research Institutes, Taiwan, R.O.C. (NHRI-EX94-9222BI, NHRI-EX96-9513SI) and the Center of Precision Medicine from the Featured Areas Research Center Program within the framework of the Higher Education Sprout Project by the Ministry of Education in Taiwan to F.-J.S.L.

## Author contributions

L.C. Chang and F.-J. S. Lee. designed the study and interpreted the results. L.C. Chang, Y.-C. Wu, and Y.-Y. Chang performed the experiments and analyzed the data. L.C. Chang and Y.-C. Wu prepared the draft of the manuscript. L.C. Chang and F.-J. S. Lee wrote and edited the manuscript.

## Conflict of interest statement

None declared.

## Notes

### Competing Interest Statement

The authors have declared no competing interest.

