## Supplemental Figures and Tables for "MAP kinase Slt2p attenuates cell wall mRNA decay by downregulating the RNA-binding protein Rbp1p in response to stress"

**Supplemental information**

**Supplemental Figure legends**

**Supplemental Figure 1. Expression levels of Rbp1p and Psp1p.**

(A) Western blotting analyzed the expression of endogenous Psp1p or exogenous expressed GFP-Psp1p in BY4741 wild-type cells, *slt2Δ* mutants and *psp1Δ* mutants. (B and C) Western blotting analyzed endogenous and exogenously expressed Rbp1p (B) and confirmed the deletion of *RBP1* (C) in YTC345 *rbp1Δ* strains.

**Supplemental Figure 2. Deletion of *RBP1* did not relieve growth perturbation from caffeine or heat shock-induced cell wall damage.**

(A and B) BY4741 wild-type, *slt2Δ*, *rbp1Δ*, or *slt2Δrbp1Δ* mutant cells were grown to log phase in rich medium, serially tenfold diluted and spotted onto rich agar medium in the presence of caffeine (4 or 15 mM) (A) and then incubated at 30 or 38°C (B) for 2 to 3 days.

**Supplemental Figure 3. Rbp1p-targeted cell wall mRNAs are stabilized in response to cell wall stress.**

YTC345 (*rpb1-1*) wild-type cells were grown in rich medium to log-phase, untreated or treated with 25 μg/ml Congo Red at permissive temperature for 2 hr followed by shifting to nonpermissive temperature to shut off transcription. Northern blot analysis, quantification and statistical analysis were performed as described in Figure 1.

**Supplemental Figure 4. The yeast two-hybrid assays of Rbp1p and Slt2p with various deletions.**

(A) Schematic representation of the Slt2p protein domain structure. C-terminal truncated variants and C-terminal nonkinase region. (B) Kinase domain (KD) alone or C-terminal nonkinase region (CF) were insufficient to interact with Rbp1p in yeast two-hybrid assays. YEM1α cells carrying LexA- and Gal4AD-based fusion constructs as indicated were assayed for β-galactosidase activity.

**Supplemental Figure 5. Congo Red treatment induces the degradation of Rbp1p but not *RBP1* transcript.**

**Supplemental Figure 6. The kinase activity of Slt2p and the upstream signal are required to rescue *slt2*Δ mutants in cell growth (A, B) and in the control of Rbp1p stability (C, D) in response to cell wall stress.**

(A) Schematic representation of the Slt2p protein domain structure with kinase activity-related mutants. (B) BY4741 wild-type carrying empty vector or *slt2Δ* expressing empty vector, full-length *SLT2*, *TYAF* or *K54R* cells were grown to log phase in synthetic selection medium, serially tenfold diluted and spotted onto synthetic selection agar medium or rich agar medium in the presence of Congo Red (10 or 25 μg/ml) and then incubated at 30 for 2 to 3 days. (C and D) Slt2p-TYAF or -K54R fails to decrease Rbp1p protein during cell wall stress. *slt2*Δ mutants expressing Slt2p Slt2p-K54R (C) or Slt2p-TYAF (D) were treated with Congo Red for the indicated times. Western blots show Rbp1p protein levels in BY4741 cells. The levels of Rbp1p were quantified by ImageJ, normalized relative to those of Act1p, and then graphed as relative levels of Rbp1p in 0 min. Standard deviations from three independent experiments are indicated.

**Supplemental Figure 7. Slt2p-dC126 shows different effectivities on death phenotype complementation of slt2Δ strain under cell wall stresses. Comparable expression levels of Slt2p, Slt2p-dC115, and Slt2p-dC126 (A), Congo Red (B), and caffeine (C) treatments.**

(A) Western blotting to show the expression level of Slt2p and its truncated variants. (B and C) BY4741 wild-type carrying empty vector or *slt2Δ* expressing empty vector, full-length *SLT2*, *dC115* or *dC126* cells were grown to log phase in synthetic selection medium, serially tenfold diluted and spotted onto synthetic selection agar medium or rich agar medium in the presence of Congo Red (10 or 25 μg/ml) (B) or caffeine (4 or 15 mM) (C) and then incubated at 30 for 2 to 3 days.

#### Supplemental Figure 1

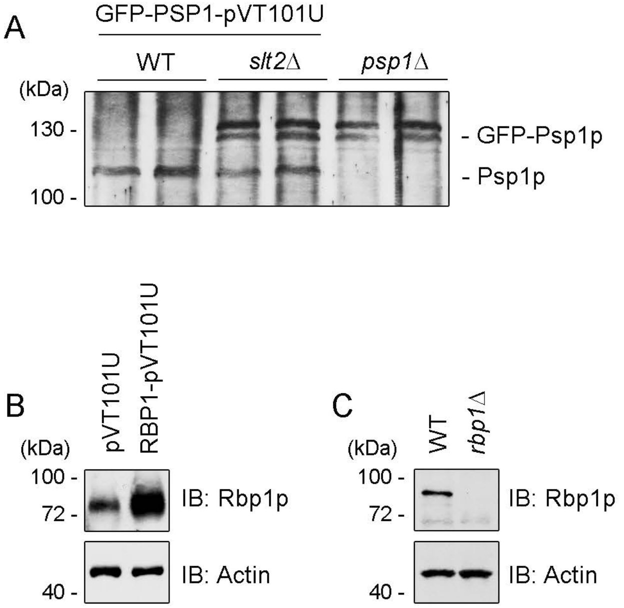

### Supplemental Figure 2

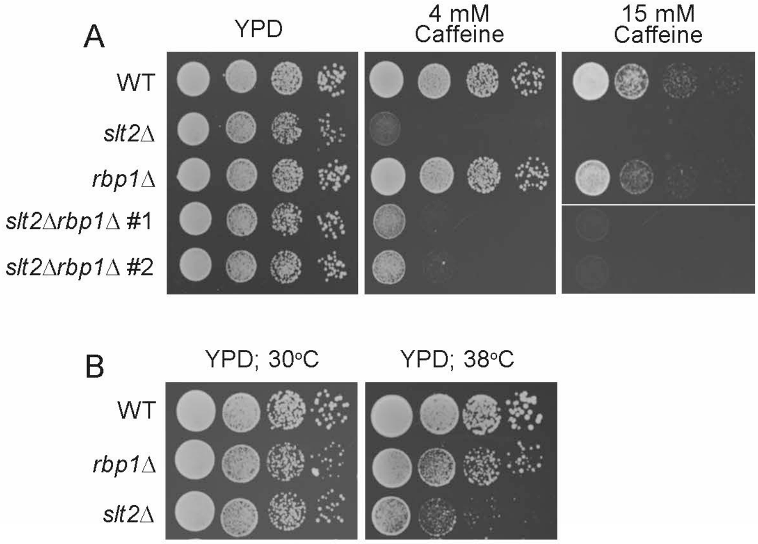

### Figure S3

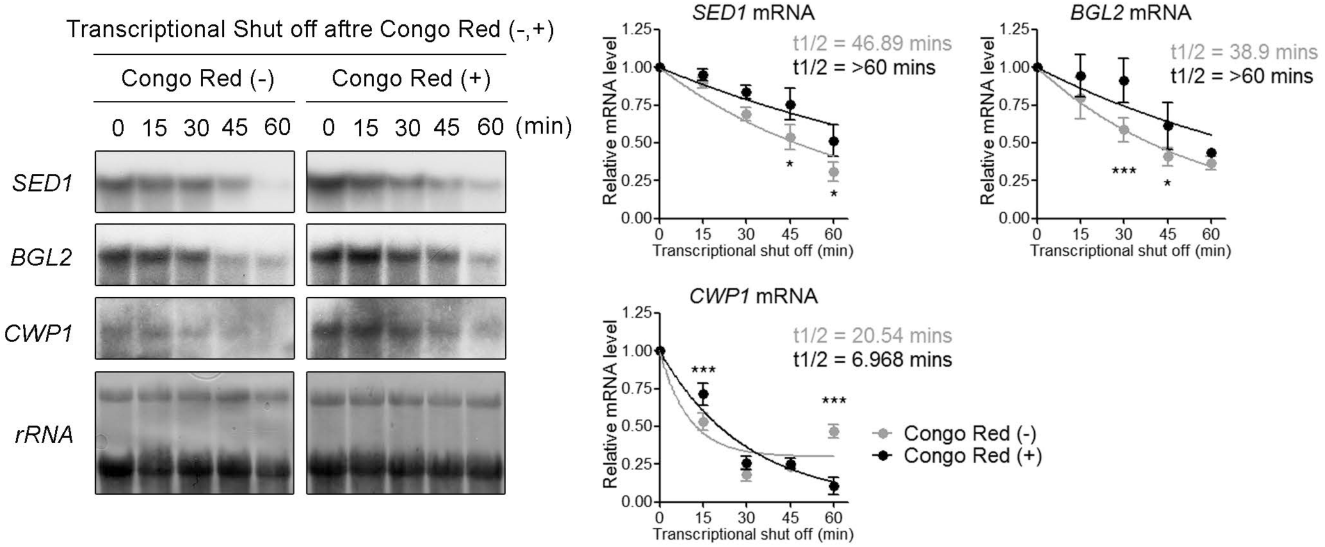

Transcriptional Shut off aftre Congo Red (-,+)
Congo Red (-)
Congo Red (+)
0 15 30 45 60 0 15 30 45 60 (min)
SED1
BGL2
CWP1
rRNA
SED1 mRNA
t1/2 = 46.89 mins
t1/2 = >60 mins
BGL2 mRNA
t1/2 = 38.9 mins
t1/2 = >60 mins
CWP1 mRNA
t1/2 = 20.54 mins
t1/2 = 6.968 mins
Relative mRNA level
Transcriptional shut off (min)
Congo Red (-)
Congo Red (+)

Supplemental Figure 4

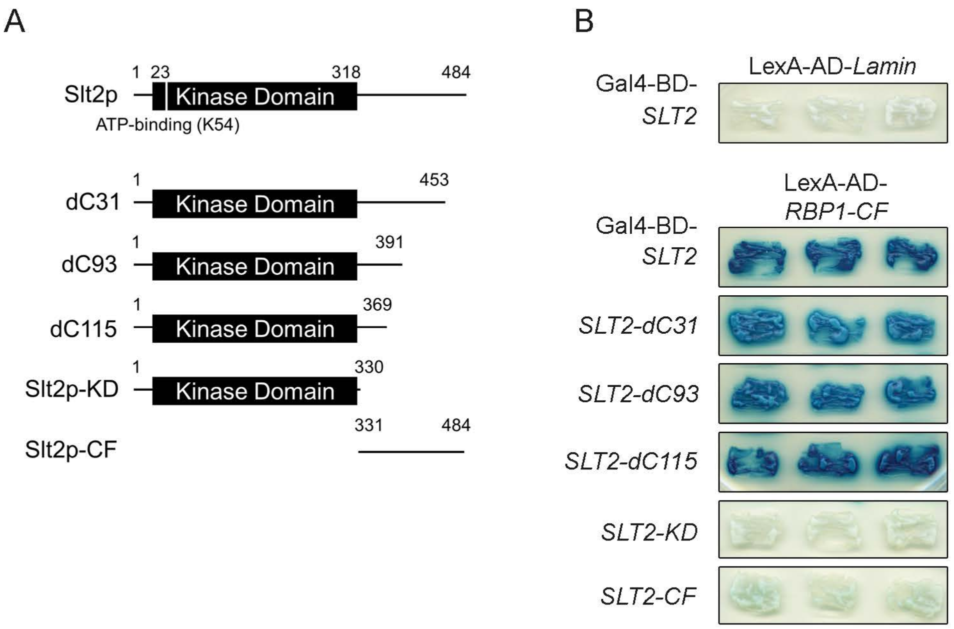

### Supplemental Figure 5

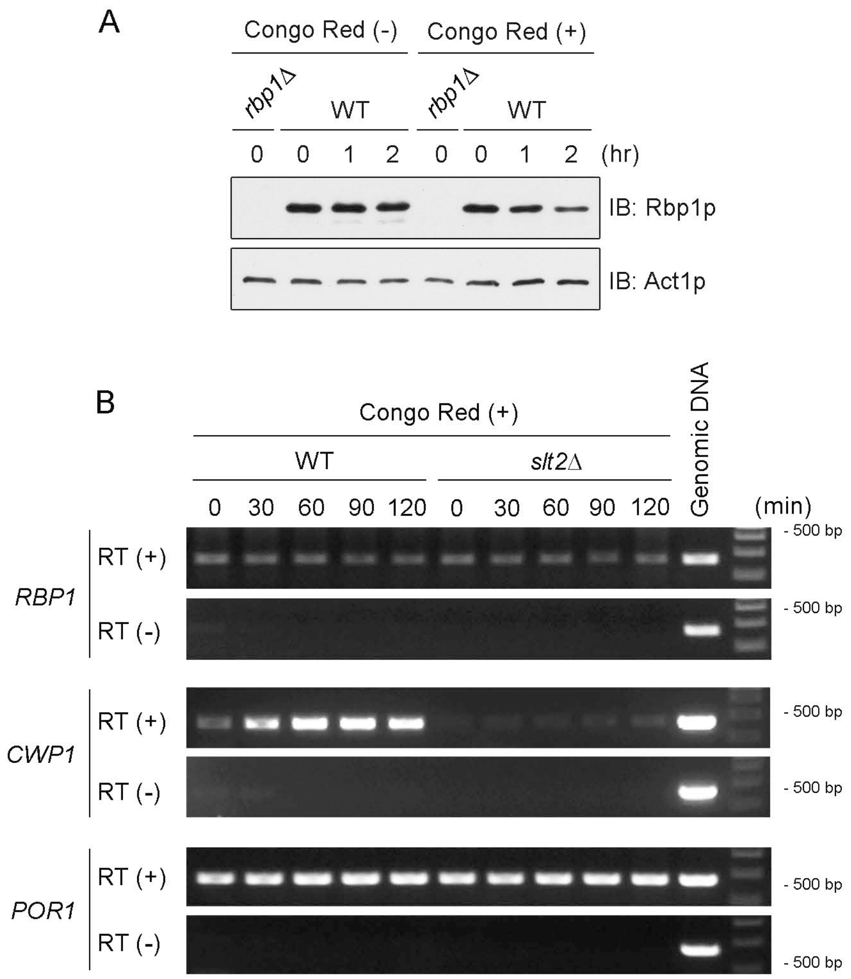

### Supplemental Figure 6

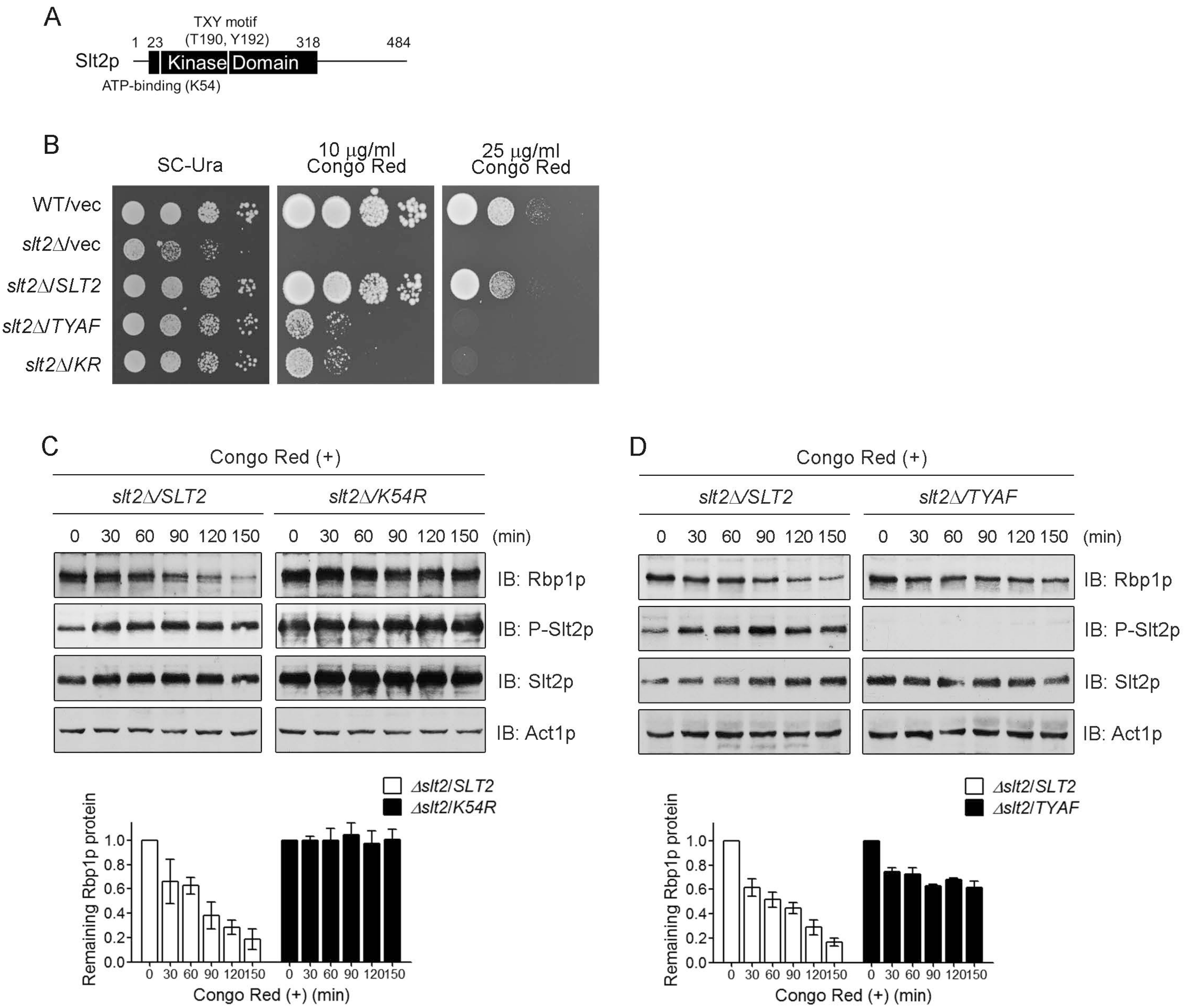

### Supplemental Figure 7

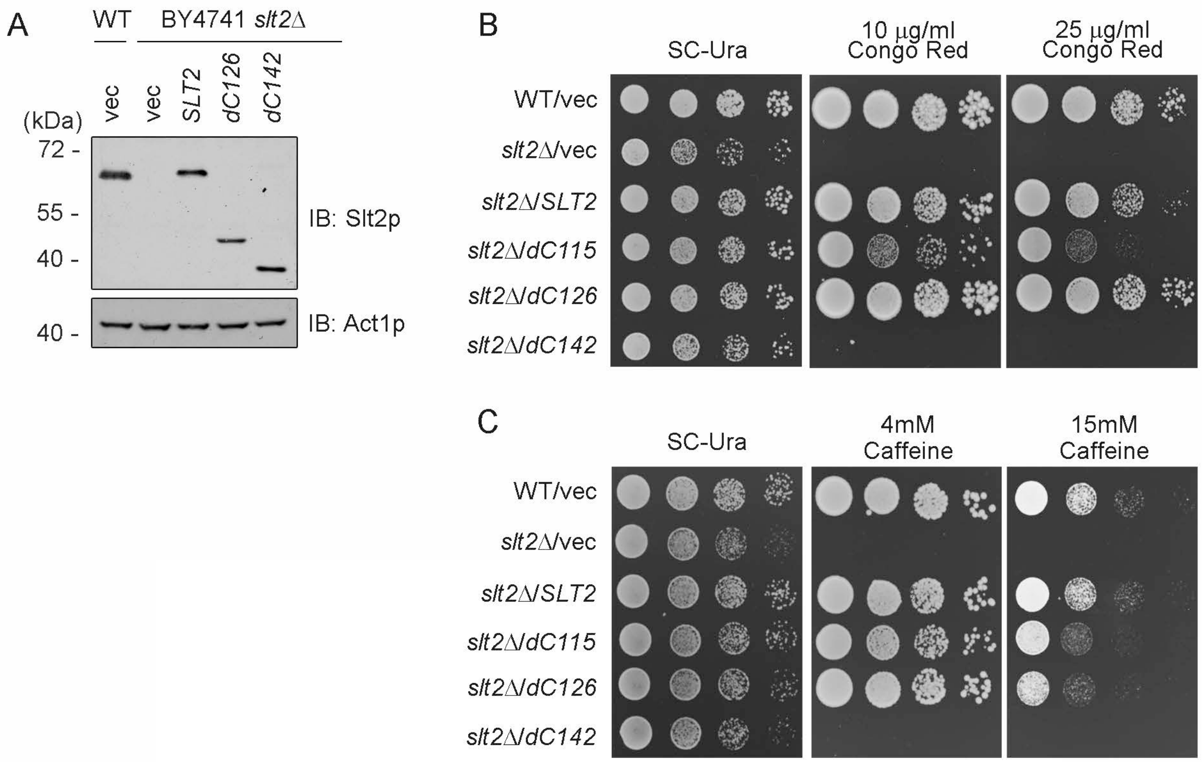

**Supplemental Table I.** Cell wall mRNAs displaying level changes in Rbp1p microarrays

| ORF | Gene | Function |
|---|---|---|
| YDR077W | *SED1* | Stress-induced GPI-cell wall protein |
| YKL096W | *CWP1* | Cell wall mannoprotein |
| YDR055W | *PST1* | GPI-cell wall protein |
| YDR261C | *EXG2* | Exo-1,3-beta-glucanase |
| YLR342W | *GSC1* | 1,3-beta-D-glucan synthase |
| YGR032W | *GSC2* | 1,3-beta-D-glucan synthase |
| YER150W | *SPI1* | GPI-cell wall protein |
| YGR189C | *CRH1* | Chitin transglycosylase |
| YNL066W | *SUN4* | Cell wall protein related to glucanases |
| YGR279C | *SCW4* | Cell wall protein related to glucanases |
| YKL096W-A | *CWP2* | Cell wall mannoprotein |
| YLR110C | *CCW12* | Cell wall mannoprotein |
| YKL163W | *PIR3* | O-glycosylated covalently-bound cell wall protein |
| YOR383C | *FIT3* | Cell wall mannoprotein |
| YNL327W | *EGT2* | GPI-cell wall endoglucanase |
| YOR247W | *SRL1* | Cell wall mannoprotein |
| YGR282C | *BGL2* | Endo-1,3-beta-glucanase |
| YJL159W | *HSP150* | O-mannosylated heat shock protein |

**Supplemental Table II.** Strains used in this study

| Strain | Genotype | Source |
|---|---|---|
| YEM1α | *MAT**a** his3 trp1 leu2 6ops-LEU2 2ops-LacZ* | |
| BY4741 | *MAT**a** his3 leu2 ura3 met15* | |
| BY4741*slt2Δ* | BY4741 except *slt2::KanMX6* | ResGen (Invitrogen) |
| BY4741*dhh1Δ* | BY4741 except *dhh1::KanMX6* | ResGen (Invitrogen) |
| BY4741*pat1Δ* | BY4741 except *pat1::KanMX6* | ResGen (Invitrogen) |
| BY4741*lsm1Δ* | BY4741 except *lsm1::KanMX6* | ResGen (Invitrogen) |
| BY4741*edc3Δ* | BY4741 except *edc3::KanMX6* | ResGen (Invitrogen) |
| BY4741*ski2Δ* | BY4741 except *ski2::KanMX6* | ResGen (Invitrogen) |
| BY4741*rbp1Δ* | BY4741 except *rbp1::KanMX6* | ResGen (Invitrogen) |
| BY4741*slt2Δrbp1Δ* | BY4741 except *slt2::KanMX6 rbp1::HisMX6* | This study |
| BY4741*slt2Δdhh1Δ* | BY4741 except *slt2::KanMX6 dhh1::HisMX6* | This study |
| BY4741*slt2Δpat1Δ* | BY4741 except *slt2::KanMX6 pat1::HisMX6* | This study |
| BY4741*slt2Δlsm1Δ* | BY4741 except *slt2::KanMX6 lsm1::HisMX6* | This study |
| BY4741*slt2Δedc3Δ* | BY4741 except *slt2::KanMX6 edc3::HisMX6* | This study |
| BY4741*slt2Δski2Δ* | BY4741 except *slt2::KanMX6 ski2::HisMX6* | This study |
| YTC345 | *MAT**a** rpb1-1ura3 leu2* | Dr. Tien-Hsien Chang |
| YTC345*slt2Δ* | YTC345 except *slt2::KanMX6* | This study |
| YTC345*slt2Δrbp1Δ* | YTC345 except *slt2::KanMX6 rbp1::URA* | This study |

**Supplemental Table III.** Plasmids used in this study

| Plasmid | Features | Source |
|---|---|---|
| pVT101U | *URA3*, 2 µm, *ADH1p* | (1) |
| pVT101U-HA-RBP1 | *URA3*, 2 µm, *ADH1p-HA-RBP1* | (2) |
| pVT101U-SLT2-HA | *URA3*, 2 µm, *ADH1p-SLT2-HA* | This study |
| pVT101U-SLT2-dC126-HA | *URA3*, 2 µm, *ADH1p-SLT2-dC126-HA* | This study |
| pVT101U-SLT2-K54R | *URA3*, 2 µm, *ADH1p-SLT2-K54R* | This study |
| pVT101U-SLT2-TYAF | *URA3*, 2 µm, *ADH1p-SLT2-TYAF* | This study |
| pVT101U-HA-SLT2 | *URA3*, 2 µm, *ADH1p-HA-SLT2* | This study |
| pVT101U-HA-ERK2 | *URA3*, 2 µm, *ADH1p-HA-ERK2* | This study |
| pVT101U-HA-ERK2-C126 | *URA3*, 2 µm, *ADH1p-HA-ERK2-C126* | This study |
| pVT101U-HA-ERK2-KD | *URA3*, 2 µm, *ADH1p-HA-ERK2-KD* | This study |
| pVT101U-HA-ERK2-dC5 | *URA3*, 2 µm, *ADH1p-HA-ERK2-dC5* | This study |
| pVT101U-HA-ERK2-dC23 | *URA3*, 2 µm, *ADH1p-HA-ERK2-dC23* | This study |
| pVT101U-HA-ERK2-dC47 | *URA3*, 2 µm, *ADH1p-HA-ERK2-dC47* | This study |
| pVT101U-ERK2-HA | *URA3*, 2 µm, *ADH1p-ERK2-HA* | This study |
| pVT101U-ERK2-dC5-HA | *URA3*, 2 µm, *ADH1p-ERK2-dC5-HA* | This study |
| pVT101U-ERK2-dC23-HA | *URA3*, 2 µm, *ADH1p-ERK2-dC23-HA* | This study |
| pVT101U-ERK2-dC47-HA | *URA3*, 2 µm, *ADH1p-ERK2-dC47-HA* | This study |
| pVT101U-HA-ERK2-dC47-HA | *URA3*, 2 µm, *ADH1p-HA-ERK2-dC47-HA* | This study |
| pEGKT-RBP1 | *URA3*, 2 µm, *GAL1/10-GST-RBP1* | Our lab |
| YCplac111-RBP1 | *LEU2*, *CEN4*, *ADH1p-RBP1* | This study |
| YCplac111-RBP1-3A | *LEU2*, *CEN4*, *ADH1p-RBP1-T189A-S552A-T637A* | This study |
| pEG202 | *HIS3*, 2 µm, *ADH1p* | (3) |
| pEG202-RBP1 | *HIS3*, 2 µm, *ADH1p-RBP1* | Our lab |
| pEG202-RBP1-NF | *HIS3*, 2 µm, *ADH1p-RBP1-NF* | Our lab |
| pEG202-RBP1-dC | *HIS3*, 2 µm, *ADH1p-RBP1-dC* | Our lab |
| pEG202-RBP1-CT | *HIS3*, 2 µm, *ADH1p-RBP1-CT* | Our lab |
| pEG202-RBP1-CF | *HIS3*, 2 µm, *ADH1p-RBP1-CF* | Our lab |
| pEG202-RBP1-dN | *HIS3*, 2 µm, *ADH1p-RBP1-dN* | Our lab |
| pEG202-LAMIN | *HIS3*, 2 µm, *ADH1p-LAMIN* | Our lab |
| pJG4-5 | *TRP1*, 2 µm, *GAL1p-HA* | (3) |
| pJG4-5-SLT2 | *TRP1*, 2 µm, *GAL1p-HA-SLT2* | This study |
| pJG4-5-SLT2-dC31 | *TRP1*, 2 µm, *GAL1p-HA-SLT2-dC31* | This study |
| pJG4-5-SLT2-dC93 | *TRP1*, 2 µm, *GAL1p-HA-SLT2-dC93* | This study |
| pJG4-5-SLT2-dC115 | *TRP1*, 2 µm, *GAL1p-HA-SLT2-dC115* | This study |
| pJG4-5-SLT2-dC126 | *TRP1*, 2 µm, *GAL1p-HA-SLT2-dC126* | This study |

| | | |
|---|---|---|
| pJG4-5-SLT2-KD | *TRP1*, 2 μm, *GAL1p-HA-SLT2-K54R* | This study |
| pJG4-5-SLT2-CF | *TRP1*, 2 μm, *GAL1p-HA-SLT2-CF* | This study |
| pJG4-5-ERK2 | *TRP1*, 2 μm, *GAL1p-HA-ERK2* | This study |
| pJG4-5-ERK2-C126 | *TRP1*, 2 μm, *GAL1p-HA-ERK2-C126* | This study |
| pJG4-5-ERK2-KD | *TRP1*, 2 μm, *GAL1p-HA-ERK2-KD* | This study |
| pJG4-5-ERK2-dC5 | *TRP1*, 2 μm, *GAL1p-HA-ERK2-dC5* | This study |
| pJG4-5-ERK2-dC23 | *TRP1*, 2 μm, *GAL1p-HA-ERK2-dC23* | This study |
| pJG4-5-ERK2-dC47 | *TRP1*, 2 μm, *GAL1p-HA-ERK2-dC47* | This study |
| pGEX-6P-1-RBP1-CF | *f1, ori, Amp, T7p-GST-RBP1-CF* | Our lab |
| pET32b-SLT2 | *f1, ori, Amp, T7p-His-SLT2* | This study |
| pET32b-SLT2-dC126 | *f1, ori, Amp, T7p-His-SLT2-dC126* | This study |
